# Elicitation of neutralizing antibody responses to HIV-1 immunization with nanoparticle vaccine platforms

**DOI:** 10.1101/2021.06.03.446978

**Authors:** Amyn A. Murji, Juliana S. Qin, Tandile Hermanus, Lynn Morris, Ivelin S. Georgiev

## Abstract

A leading strategy for developing a prophylactic HIV-1 vaccine is the elicitation of antibodies that can neutralize a large fraction of circulating HIV-1 variants. However, a major challenge that has limited the effectiveness of current vaccine candidates is the extensive global diversity of HIV-1 Env, the sole target for HIV-neutralizing antibodies. To address this challenge, a variety of strategies to incorporate Env diversity into the vaccine formulation have been proposed. Here, we assessed the potential of two such strategies that utilize a nanoparticle-based vaccine platform to elicit broadly neutralizing antibody responses. The nanoparticle immunogens developed here consisted of different formulations of Envs from strains BG505 (clade A) and CZA97 (clade C), attached to the N-termini of bacterial ferritin. Single-antigen nanoparticle cocktails as well as mosaic nanoparticles bearing both Env trimers elicited high antibody titers in mice and guinea pigs.

Furthermore, serum from guinea pigs immunized with nanoparticle immunogens achieved autologous, and in some cases heterologous, tier 2 neutralization, although significant differences between mosaic and single-antigen nanoparticles were not observed. These results provide insights into the ability of different vaccine strategies for incorporating Env sequence diversity to elicit neutralizing antibodies, with implications for the development of broadly protective HIV-1 vaccines.

## INTRODUCTION

The extreme genetic diversity of circulating HIV-1 strains has been a barrier to the development of an efficacious vaccine. This is due, in part, to variations in the envelope trimer protein (Env) of HIV-1 (Cuevas et al., 2015; Williamson, 2003). Env mediates virus-host cell fusion and is also the sole target of neutralizing antibodies (Blumenthal et al., 2012). This diversity therefore underscores the need for immunogens that can generate a protective antibody response capable of recognizing a multitude of Envs. Multiple, soluble Env trimers have been used in sequence and/or simultaneously in immunizations to limited effect (Escolano et al., 2016; Klasse et al.; Torrents de la Peña et al., 2018). In addition to soluble trimers, multimerizing Env antigens on the surface of nanoparticles have also been proposed to further improve on the elicited antibody responses (Sliepen et al., 2015; Yassine et al., 2015).

Heterologous nanoparticles are capable of mimicking repetitive multimeric patterns that are recognized by the immune system, engage with antigen presenting cells, and traffic directly to the lymphatic system (Martin et al., 2020; Tokatlian, 2019). The arrangement of antigens on a nanoparticle can effectively cross-link B-cell receptors, which improves antibody-dependent immune responses (Bachmann and Zinkernagel, 1997; He et al., 2016). Spontaneously self-assembling protein nanoparticles have been particularly efficient, as their expression in culture circumvents additional steps of nanoparticle-antigen linkage (Kanekiyo et al., 2013; Li et al., 2006; Sliepen et al., 2015). Different-sized protein nanoparticles have been tested against a range of disease states, exhibiting their versatility as a scaffold for antigen presentation (López-Sagaseta et al., 2016).

*Heliobacter pylori* ferritin in particular has been used as a platform for many viral antigens, including HIV-1 and influenza (Kanekiyo et al., 2013, 2015; Li et al., 2019; Sliepen et al., 2015; Swanson et al., 2020). These particles are comprised of 24 identical monomers that spontaneously self-assemble into a ∼10 nm particle that includes eight three-fold axes of symmetry, making them an optimal candidate for expressing trimeric antigens like the HIV-1 Env. Genetically fusing diverse Env protomers to the N-terminus of self-assembling protein nanoparticles like ferritin also allows individual nanoparticles to display multiple trimers; here, we refer to such particles as “mosaic” nanoparticle immunogens (Kanekiyo et al., 2019). The mosaic nanoparticle strategy aims at generating a cross-reactive B-cell response capable of recognizing the different antigens on the surface of the nanoparticle. While mosaic nanoparticles have been successfully developed for various targets including influenza and coronavirus (Cohen et al., 2021; Kanekiyo et al., 2019), in the case of HIV-1, it is currently not well-understood if these particles outperform cocktails of single-antigen nanoparticles or other strategies for incorporating Env diversity. To address this question, we performed immunization experiments with the goal of comparing the elicited antibody responses to mosaic nanoparticles vs. alternative strategies, with each strategy using the same two underlying HIV-1 strains. Our results indicate that mosaic nanoparticles can elicit autologous, and in some cases heterologous, HIV-1 Tier 2 neutralizing antibody responses, although significant differences with single-antigen nanoparticles were not observed for the two specific strains tested here. Overall, our results provide insights into the ability of different vaccine strategies for incorporating Env sequence diversity to elicit neutralizing antibody responses, with implications for the development of broadly protective HIV-1 vaccines.

## Results

### Design and characterization of HIV-1 nanoparticle immunogens

BG505 and CZA97 HIV-1 Env trimers were designed as single chain variants (referred to as scBG505 and scCZA97) with a non-cleavable linker between gp120 and gp41 (Georgiev et al., 2015). These trimers also incorporated SOSIP stabilizing mutations and truncation at residue 664, based on HXB2 numbering (Korber et al.; Sanders et al., 2013). We also designed protein nanoparticles by genetically fusing either scBG505 or scCZA97 to the N-termini of *Heliobacter pylori* ferritin, separated by a 3-residue linker (Sliepen et al., 2015). The expression of the scBG505-ferritin plasmid or the scCZA97-ferritin plasmid in HEK293F cells resulted in particles multimerized with each respective trimer, as reported previously (Sliepen et al., 2015). We will refer to these particles as single-antigen nanoparticle immunogens (Figure 1A). In order to multimerize both scEnvs on the same particle, we cotransfected cells with both plasmids, generating mosaic nanoparticles (Figure 1B). Size exclusion chromatography profiles were used to isolate particles of the correct size (Figure S1). Nanoparticle formation was also confirmed by negative-stain EM, which identified particles approximately 40 nm wide with trimer spikes as expected and previously reported (Sliepen et al., 2015)(Figure 1C).

**Figure 1.**
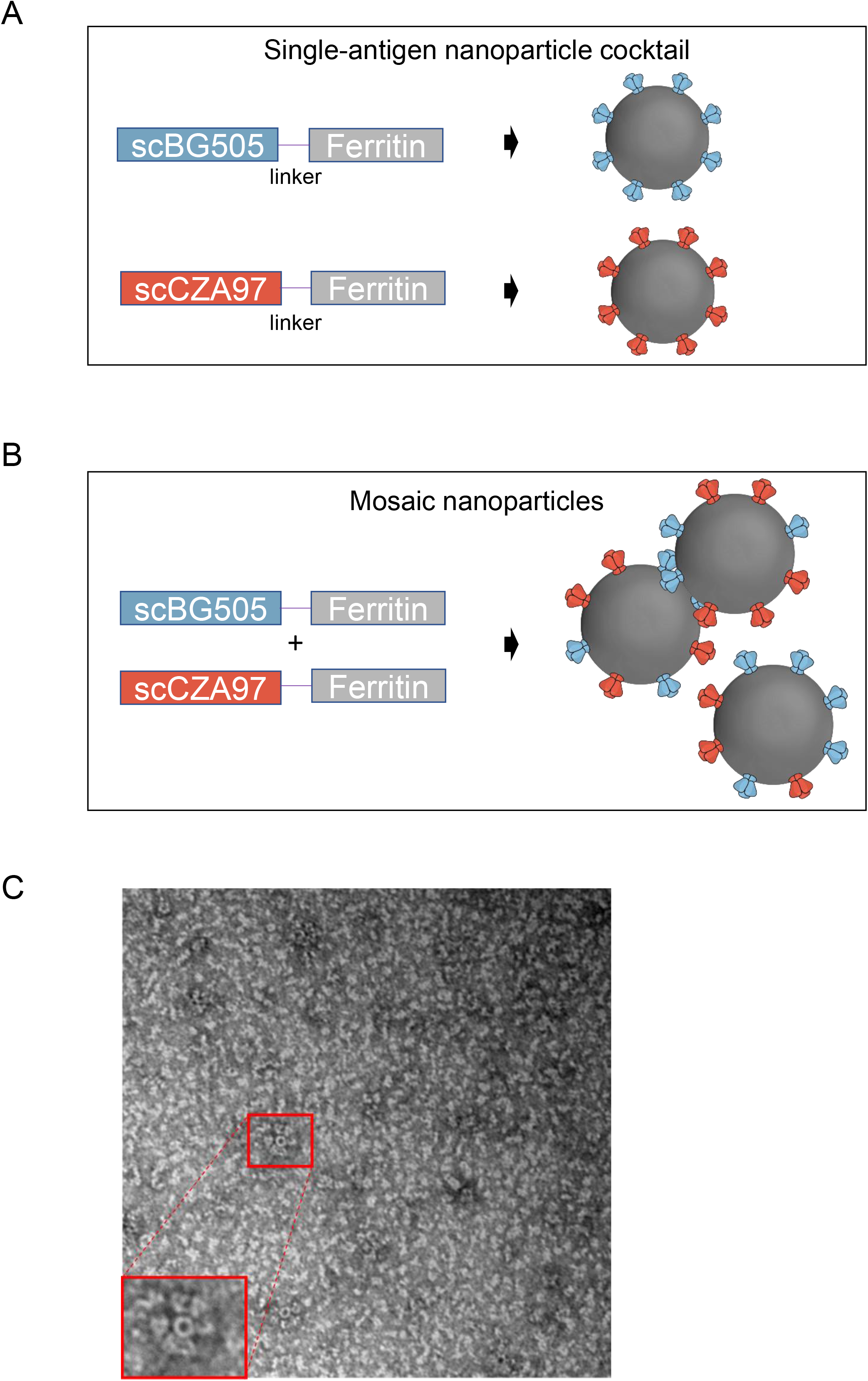
Nanoparticle immunogens. Schematic depicting the **A**. Single-antigen nanoparticle immunogens and **B**. Mosaic nanoparticles. Blue and red bars depict the single-chain protomer of envelope proteins from BG505 and CZA97, respectively. The grey bar is representative of one protomer of ferritin. **C**. Representative EM image showing nanoparticle formation.

### Single-antigen and mosaic HIV-1 nanoparticle immunogens elicit comparable responses in mice

To determine the immunogenicity of the single-antigen and mosaic nanoparticles, we first immunized mice with either (•) a cocktail of scBG505-ferritin and scCZA97-ferritin single-antigen nanoparticles, and (•) mosaic nanoparticles expressed from the cotransfection of scBG505-ferritin and scCZA97-ferritin. Two additional control groups were also added: (•) scBG505 trimer and (•) a cocktail of scBG505 and scCZA97 trimers. Mice were boosted four and eight weeks post-prime with the same immunogens, for a total of three immunizations per group. Two weeks after the final boost, mice were exsanguinated (Figure 2A). Serum from mice immunized solely with scBG505 trimer bound scBG505 and scCZA97. While not statistically significant (p=0.222, Mann-Whitney Test), titers to scCZA97 appeared to be lower compared to titers to scBG505 (geometric mean AUC values 7.2±1.5 and 3.6±2.3 for scBG505 and scCZA97, respectively). In contrast, mice immunized with cocktails of trimers (geometric mean AUC values 8.1±1.2, 8.5±1.1), cocktails of single-antigen nanoparticles (8.1±1.4, 6.9±1.3), and mosaic nanoparticles (10.1±1.1, 8.5±1.0), each elicited high titers to both scBG505 and scCZA97 proteins, respectively (Figure 2B). Overall, no significant differences in immunogenicity were observed between the four groups (Figure 2C).

**Figure 2.**
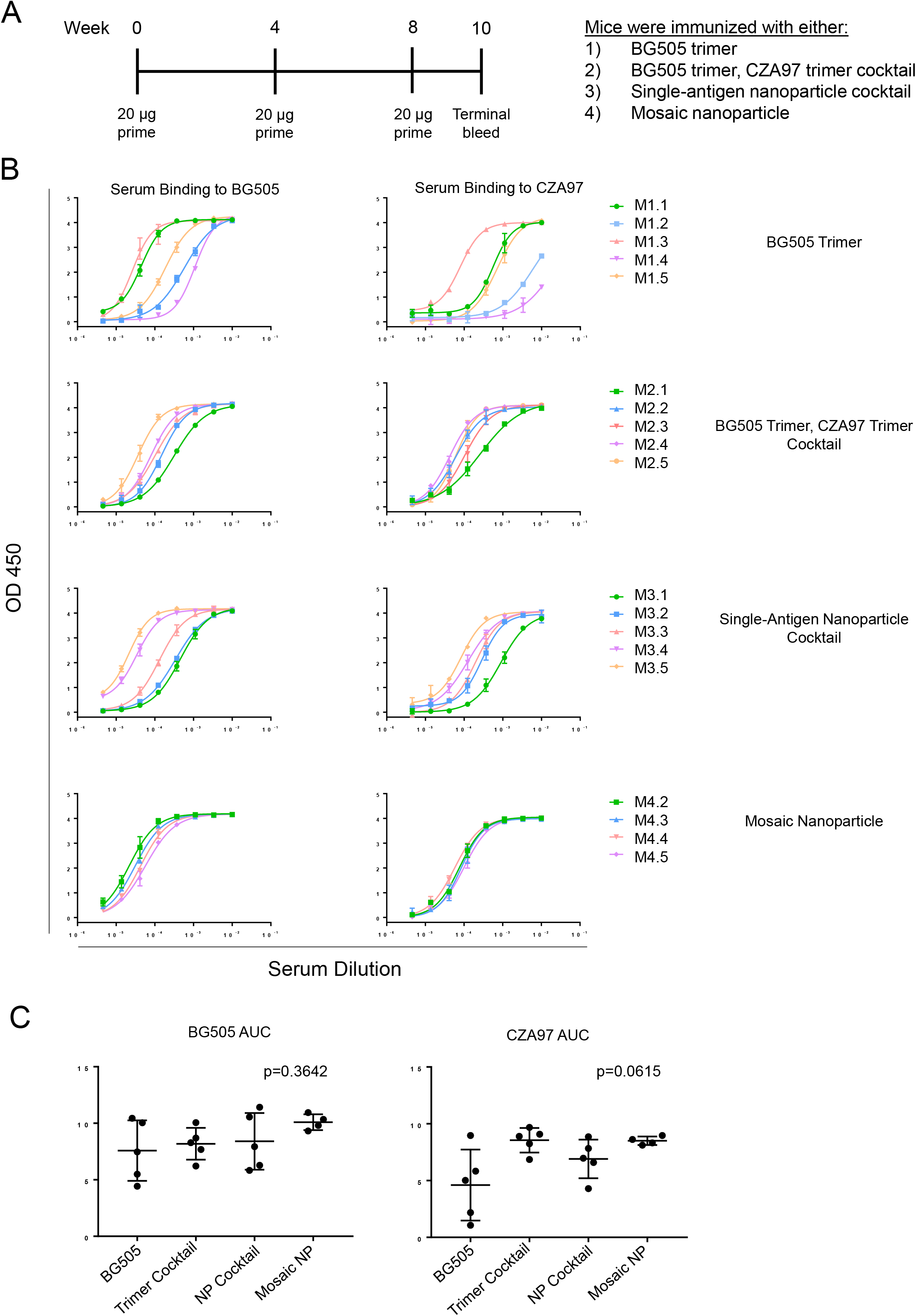
Nanoparticle immunogens elicit strong antibody responses in mice. **A**. Mouse immunization scheme. Mice (n=5) were primed at week zero and boosted every 4 weeks. Terminal bleeds were performed 2 weeks after the second (final) boost. **B**. Mice (n=5) were immunized with one of the following: BG505 trimer (M1.1-1.5), cocktail of BG505 and CZA97 trimers (M2.1-2.5), the single-antigen nanoparticle cocktail (M3.1-M3.5), or mosaic nanoparticles (M4.2-4.4). Data for M4.1 are not available due to insufficient volume of serum. Each mouse is color-coded. Curves refer to binding to BG505 (left) or CZA97 (right) trimer. **C**. AUC values from ELISA curves are shown for each group in (B). P values were calculated using Kruskal-Wallis tests and were adjusted using Dunn’s multiple comparisons test.

### HIV-1 neutralization by vaccine-elicited antibody responses in mice

Given the robust titers of serum binding to vaccine-matched (autologous) Envs, we then asked whether differences could be observed in the breadth or potency of serum neutralization. Though serum quantities were limited from some mice, we observed autologous neutralization in one mouse solely immunized with BG505. No other vaccinated mice displayed neutralization of the BG505 or CZA97 autologous strains (Figure 3).

**Figure 3.**
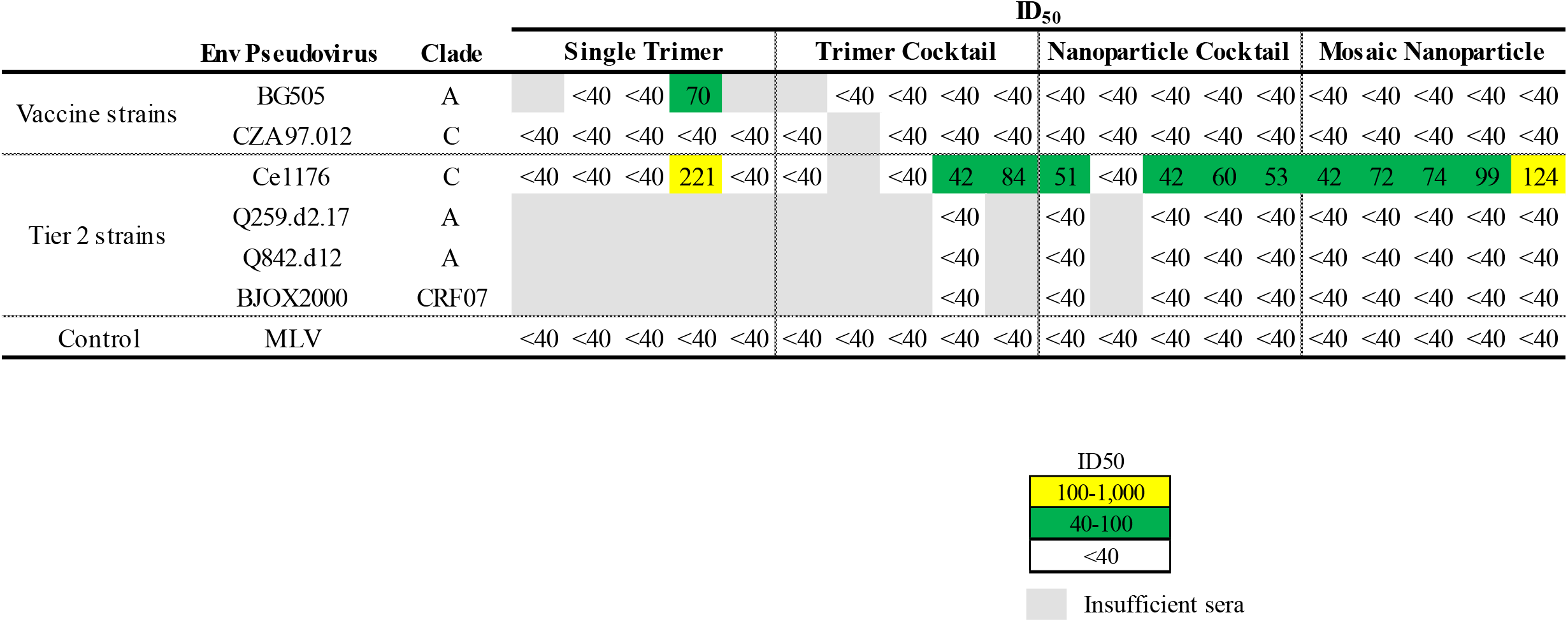
Nanoparticle immunogens elicit neutralizing antibody responses in mice. Plasma samples collected from mice (n = 20) immunized with BG505 trimer, cocktail of BG505 and CZA97 trimers, monovalent nanoparticle cocktail, or mosaic nanoparticles were tested for neutralization to vaccine-matched and Tier 2 pseudoviruses. Mouse sera neutralization titers were reported as plasma dilution required to inhibit 50% of virus infection (ID_50_). The magnitude of neutralization responses (ID_50_) is indicated by the color, from moderate potency (yellow) to lower potency (green) and no activity (white). Gray cells denote insufficient sera for testing.

Intriguingly, serum from one mouse (20%) in the single trimer group, two (40%) from the trimer cocktail group, two (40%) from the nanoparticle cocktail group and four (80%) from the mosaic nanoparticle group neutralized a third, heterologous virus from Clade C, Ce1176, but no other Tier 2 viruses (Figure 3).

### Serum from guinea pigs immunized with cocktails of nanoparticles and cotransfected nanoparticles recognize heterologous gp140 Envs

Because elicitation of broadly HIV-neutralizing antibodies has historically been a challenge in mouse models (Hu et al., 2015), we also immunized guinea pigs after confirming immunogenicity of our two types of nanoparticle immunogens. Guinea pigs have generally shown greater ability to elicit HIV-1 bNAbs and possess more blood volume, allowing for testing larger virus neutralization panels (Bricault et al., 2019; Xu et al., 2018). Guinea pigs were immunized four times, three weeks apart with either (•) cocktails of scBG505-ferritin and scCZA97-ferritin single-antigen nanoparticles or (•) cotransfected scBG505-ferritin and scCZA97-ferritin mosaic nanoparticles (Figure 4A). Serum from terminal bleeds were tested for binding against vaccine-matched trimers, with both groups generally eliciting high titers against vaccine-matched trimers, with the exception of one animal in the single-antigen cocktail group (Figure 4B, S2A). To determine whether the two groups resulted in different breadth of HIV-1 Env variant recognition, we tested the vaccine sera against a panel of diverse gp140 Envs (Figure 4C, S2B,C). A similar pattern of antigen binding was observed for both groups against the different antigens in the panel (again, with the exception of the same single animal from the single-antigen cocktail group). Together, these results suggest that both nanoparticle immunogen formulations can elicit antibodies capable of recognizing Envs from multiple, diverse clades.

**Figure 4.**
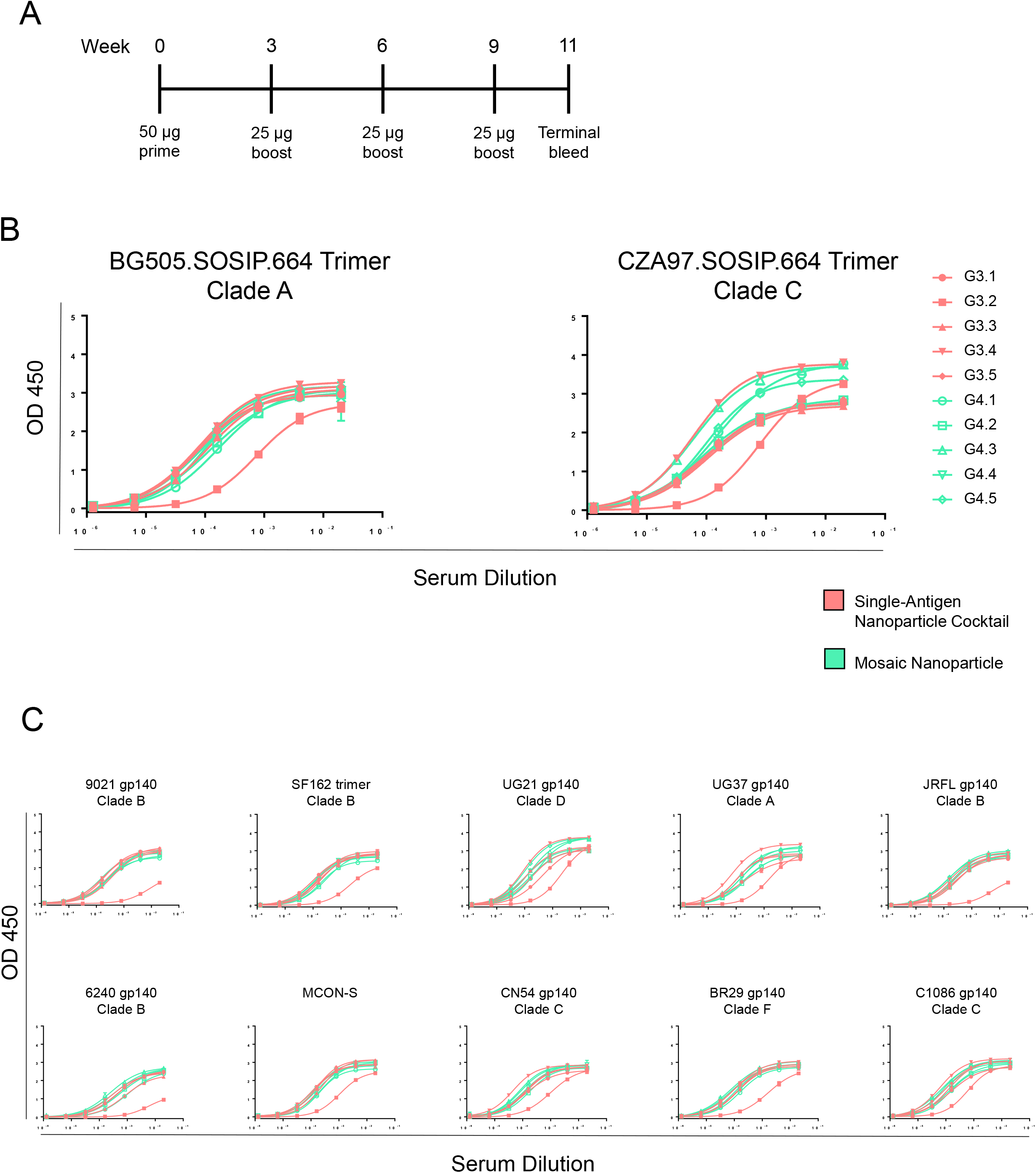
Guinea pigs immunized with nanoparticles show breadth of reactivity against diverse HIV-1 Envs. **A**. Guinea pig immunization scheme. Guinea pigs were primed at day zero and boosted every 3 weeks. Terminal bleeds were performed 2 weeks after the final boost. **B**. ELISA binding curves against vaccine-matched Envs. Guinea pigs (n=5) were immunized with either the single-antigen nanoparticle cocktail (G3.1-3.5) or mosaic nanoparticles (G4.1-4.5). Each guinea pig is color-coded by immunization group. Curves refer to binding to BG505 (left) or CZA97 (right) trimer. **C**. ELISA binding curves to gp140 Env proteins from diverse clades.

### Single-antigen and mosaic HIV-1 nanoparticle immunogens elicit autologous and heterologous virus neutralizing antibodies

Sera from guinea pigs from both immunization groups potently neutralized tier 1A and tier 1B viruses (Figure 5), with the exception of one guinea pig from the single-antigen nanoparticle cocktail group that had displayed consistently lower binding to Envs (Figure 4B,C). We next assessed whether guinea pig sera could neutralize the two vaccine-matched (autologous) viruses used for the immunizations. In guinea pigs immunized with the mosaic nanoparticle, 3/5 (60%) neutralized BG505 and 4/5 (80%) neutralized CZA97. Autologous responses to BG505 and CZA97 viruses were observed in 3/5 (60%) guinea pigs immunized with the cocktail.

**Figure 5.**
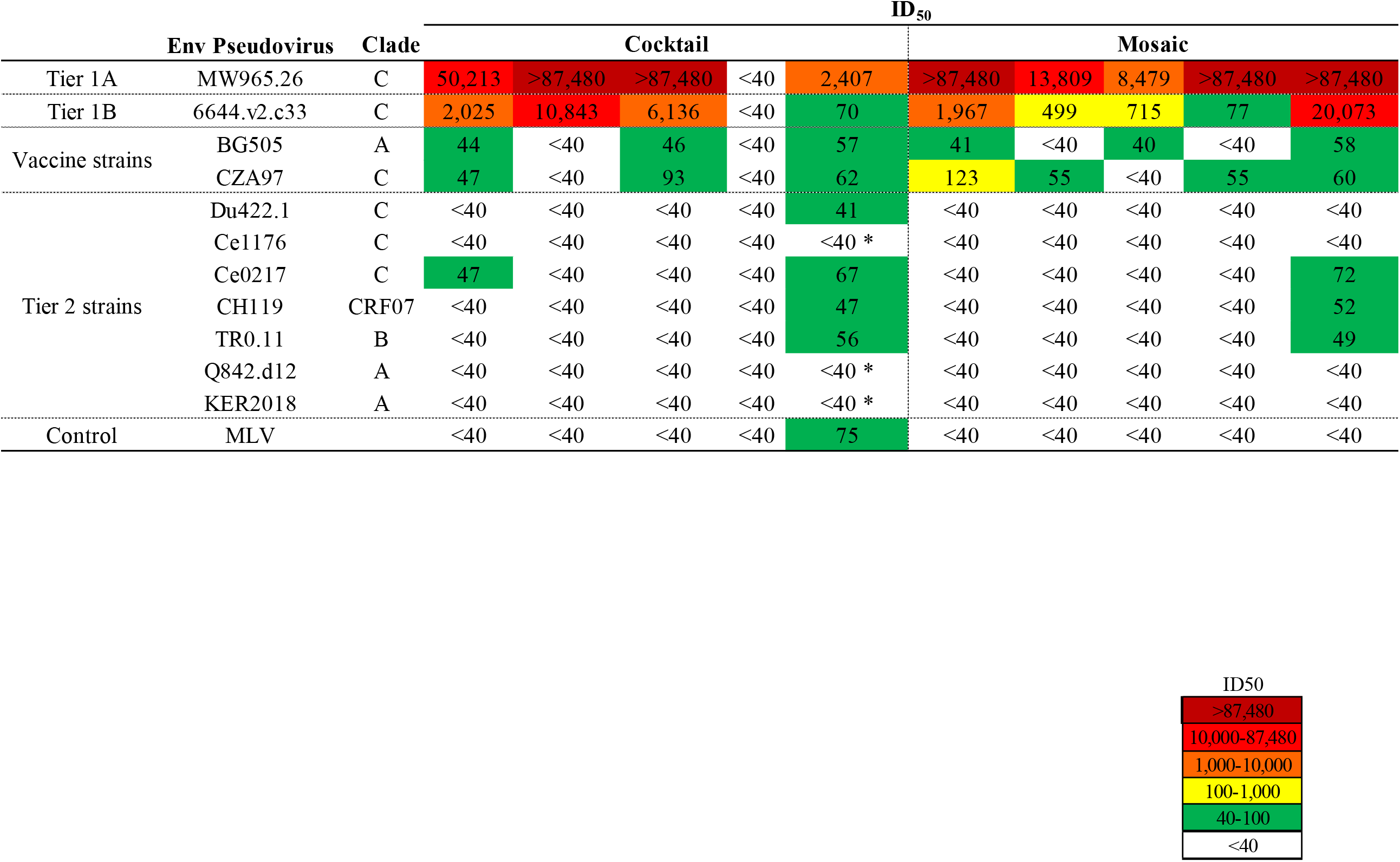
Nanoparticle immunogens elicit vaccine-matched and heterologous neutralization in guinea pigs. Plasma samples collected from guinea pigs (n = 10) immunized with either single-antigen nanoparticle cocktails or as mosaic nanoparticles bearing both BG505 and CZA97 were tested for neutralization to vaccine-matched, Tier 1, and Tier 2 pseudoviruses. The magnitude of neutralization responses (ID_50_) is indicated by the color, from higher potency (dark red) to lower potency (green) and no activity (white). Asterisks indicates titers just below 50% cut-off.

One animal from the mosaic nanoparticle group exhibited some neutralization breadth by neutralizing three of seven heterologous tier 2 viruses from diverse clades at low titers (Figure 5). Low titer neutralization of a single tier 2 heterologous virus was also seen in one animal from the single-antigen nanoparticle group. Another animal from the single-antigen nanoparticle group showed neutralization of all 7 heterologous tier 2 viruses (3 were just below the cut-off) but also neutralized the MLV control, indicating potential toxicity issues in this serum sample (Figure 5).

## DISCUSSION

Since antibodies that neutralize one HIV-1 virus do not necessarily confer protection against other circulating strains, heterologous neutralization is an important goal of HIV-1 vaccinology. It is therefore important to design immunogens capable of eliciting such responses. In particular, nanoparticle immunogens have emerged as viable alternatives to soluble trimers due to size, geometry, and the capacity to uniformly display antigens on the surface. Here, we compared cocktails of single-antigen nanoparticles and mosaic nanoparticles simultaneously displaying both Env proteins, showing immunogenicity in mice and both vaccine-matched and heterologous neutralization in guinea pigs.

Our efforts provide preliminary justification for immunizing with multiple envelope proteins from different clades expressed onto nanoparticles. While no autologous responses were observed in mice, as with many other studies (Hu et al., 2015; Sliepen et al., 2015), this animal model has historically been useful to determine immunogenicity. Neutralization of the heterologous Ce1176 virus in mice was unexpected, considering the lack of observed autologous responses in the same mice or analogous Ce1176 neutralization in guinea pigs, although Ce1176 neutralization has been observed in other vaccine studies as well (Pauthner et al., 2017). These results potentially highlight the notable differences in the ability of the immune systems of different animal models to elicit HIV-neutralizing antibody responses.

The lack of significant differences in elicited neutralization breadth between single-antigen and mosaic nanoparticles is notable, given the established advantages of mosaic nanoparticles for a number of other pathogens (Cohen et al., 2021; Kanekiyo et al., 2019). However, it is possible that optimizing additional variables, such as through specific selection of Env antigens based on bNAb epitope availability, glycan coverage, and sequence diversity may be of critical significance to fully harness the power of mosaic nanoparticles for HIV-1 vaccine design. Nevertheless, the autologous and, in some cases, heterologous tier 2 neutralization in guinea pigs observed in this study highlights the potential of multimerizing diverse Env variants on self-assembling protein nanoparticles as a viable direction for the design of a broadly protective HIV-1 vaccine.

## Acknowledgements

We thank Dr. Carol Crowther for project management and members of the Georgiev lab for comments on the manuscript. This work was supported in part by institutional funding from Vanderbilt University Medical Center and: (I.S.G.) NIH R21 AI47768; (A.A.M) NIAID T32AI112541; (L.M.) the Medical Research Council of South Africa, and NIAID U19 AI51794 (CAPRISA) and 1U01AI136677.

## Author Contributions

Conceptualization, Ivelin Georgiev; Data curation, Amyn Murji and Tandile Hermanus; Funding acquisition, Ivelin Georgiev; Investigation, Amyn Murji, Juliana Qin and Tandile Hermanus; Methodology, Ivelin Georgiev; Project administration, Ivelin Georgiev and Lynn Morris; Resources, Ivelin Georgiev and Lynn Morris; Supervision, Lynn Morris; Validation, Amyn Murji; Visualization, Amyn Murji and Juliana Qin; Writing – original draft, Amyn Murji and Ivelin Georgiev; Writing – review & editing, Amyn Murji, Ivelin Georgiev, Juliana Qin and Lynn Morris.

## Materials and Methods

### Reagents

The following reagents were obtained from the AIDS Research and Reference Reagent Program, Division of AIDS (DAIDS), National Institute of Allergy and Infectious Diseases (NIAID), National Institutes of Health (NIH): Anti-HIV-1 gp120 Monoclonal (VRC01), from Dr. John Mascola (cat# 12033) (Wu et al., 2010). The following reagents were obtained through the NIH HIV Reagent Program, Division of AIDS, NIAID, NIH: Human Immunodeficiency Virus 1 (HIV-1) JRFL gp140 Recombinant Protein (B.JRFL gp140CF), ARP-12573, contributed by Drs. Barton F. Haynes and Hua-Xin Liao; Human Immunodeficiency Virus 1 (HIV-1) gp140 Recombinant Protein (B.9021 gp140C), ARP-12575, contributed by Drs. Barton F. Haynes and Hua-Xin Liao; Human Immunodeficiency Virus 1 (HIV-1) gp140 Recombinant Protein (C.1086 gp140C), ARP-12581, contributed by Drs. Barton F. Haynes and Hua-Xin Liao; Human Immunodeficiency Virus 1 (HIV-1) gp100 Recombinant Protein (B.6240 gp140C), ARP-12572, contributed by Drs. Barton F. Haynes and Hua-Xin Liao; Human Immunodeficiency Virus Type 1 BR029 gp140 Protein, Recombinant from CHO Cells, ARP-12066, contributed by DAIDS/NIAID; produced by Polymun Scientific; Human Immunodeficiency Virus Type 1 UG21 gp140 Protein, Recombinant from CHO Cells, ARP-12065, contributed by DAIDS/NIH (Polymun Scientific, Inc); Human Immunodeficiency Virus Type 1 UG037 gp140 Protein, Recombinant from CHO Cells, ARP-12063, contributed by DAIDS/NIAID (Polymun Scientific, Inc); Human Immunodeficiency Virus Type 1 CN54 gp140 Protein, Recombinant from CHO Cells, ARP-12064, contributed by DAIDS/NIAID; produced by Polymun Scientific, Inc; Human Immunodeficiency Virus Type 1 SF162 gp140 Trimer Protein, Recombinant from HEK293T Cells, ARP-12026, contributed by Dr. Leo Stamatatos.

### Antigen expression and purification

Each of the trimers were expressed in FreeStyle 293F mammalian cells (ThermoFisher) by transfecting plasmids corresponding to either BG505.SOSIP.664sc or CZA97.SOSIP.664sc using polyethylenimine (PEI) transfection reagent and cultured for 5-7 days. These cells were cultured at 37°C with 8% CO_2_ saturation and shaking. After transfection and 5-7 days of culture, cell cultures were centrifuged at 6000 rpm for 20 minutes.

Supernatant was 0.45 µm filtered with PES membrane Nalgene Rapid Flow Disposable Filter Units. Filtered supernatant was run over a column containing *Galanthus nivalis* snow-drop lectin that had been equilibrated with PBS. The column was washed with PBS, and proteins were eluted with 30 mL of 1 M methyl-α-D-mannopyranoside. The protein elution was buffer exchanged three times into PBS and concentrated using 10 kDa or 30kDa Amicon Ultra centrifugal filter units. Concentrated protein was run on a Superose 6 Increase 10/300 GL or Superdex 200 Increase 10/300 GL on the AKTA FPLC system, and fractions were collected on an F9-R fraction collector.

CZA97-ferritin and BG505-ferritin were expressed as above. Cotransfected nanoparticles were expressed by transfecting equal parts of CZA97-ferritin and BG505-ferritin DNA as described in (Kanekiyo et al., 2019) and were purified as above but using a Sephacryl S-500 or Superose 6 sizing column.

### Enzyme linked immunosorbent assay (ELISA)

For gp140 ELISAs, soluble proteins (Aids Reagent Program) protein were plated at 2 μg/ml overnight at 4°C. The next day, plates were washed three times with PBS supplemented with 0.05% Tween20 (PBS-T) and coated with 5% milk powder in PBS-T. Plates were incubated for one hour at room temperature and then washed three times with PBS-T. Guinea pig serum were diluted in 1% milk in PBS-T, starting at a 1:50 dilution and was subsequently serially diluted 1:5 and then added to the plate. The plates were incubated at room temperature for one hour and then washed three times in PBS-T. The secondary antibody, goat anti-guinea pig IgG conjugated to peroxidase (abcam), was added at 1:25,000 dilution in 1% milk in PBS-T to the plates, which were incubated for one hour at room temperature. Plates were washed three times with PBS-T and then developed by adding TMB substrate to each well. The plates were incubated at room temperature for 10 minutes, and then 1 N sulfuric acid was added to stop the reaction. Plates were read at 450 nm. Due to limited amounts of recovered sera, we were unable to perform ELISAs against scEnvs for one mouse in the group that received the mosaic immunogen Areas under the ELISA binding curves (AUC) were determined with GraphPad Prism 8.0.0.

### Mouse Immunizations

The study was conducted with approval by the Institutional Review Board of Van-derbilt University (IACUC Approved Protocol Number M1700115). 5 Female BALB/c mice 4-6 weeks old (Charles River) were used per group in our immunization studies. Mouse studies included 4 groups for a total number of 20 mice.

Mice either received BG505 tri-mer, BG505 and CZA97 trimer cocktail, BG505-ferritin and CZA97-ferritin cocktail, or BG505 and CZA97 both on ferritin. A prime dose of 20µg of antigen in TiterMax Gold (Sigma-Aldrich) was administered intramuscularly in the hind leg(s). Boosts were per-formed at 4 and 8 weeks post prime. Pre-immune blood was collected from the subman-dibular vein before the start of the study and two weeks after each immunization. Two weeks after the final boost, blood was collected from both a submandibular bleed as well as cardiac puncture.

### Guinea Pig Immunizations

The study was conducted according to the guidelines of Cocalico Biologicals, Inc. Animal Care and Use Committee (The IACUC Approved Protocol Number: 181024CBISTD) for the guinea pig studies. Cocalico Biologics administered vaccines by first priming with 50µg of either the single-antigen cocktail of nanoparticles or mosaic nanoparticles in TiterMax Gold (Sigma-Aldrich) and boosting with 25µg of either for-mu-lation every 3 weeks for a total of 3 boosts after prime. Immunizations were adminis-tered at multiple sites subcutaneously along the back and intramuscularly in the hind limbs as the routes of injection. No more than 0.1ml of the emulsion was used per site. Terminal bleeds were collected by Cocalico after guinea pigs were fully anesthetized.

### TZM-bl Neutralization Assays

Serum neutralization was assessed using the TZM-bl assay as described (Sarzotti-Kelsoe et al.,2014). This standardized assay measures antibody-mediated inhibition of infection of TZM-bl cells by molecularly cloned Env-pseudoviruses. Viruses that are highly sensitive to neutralization (Tier 1) and/or those representing circulating strains that are moderately sensitive (Tier 2) were included. Murine leukemia virus (MLV) was included as an HIV-specificity control. Neutralization was measured as a reduction in luciferase gene expression after a single round infection of TZM--bl cells in a 96 well-plate. Results are presented as the serum/plasma dilution required to inhibit 50% of virus infection (ID_50_).

### Statistical Analysis

GraphPad Prism 8.0.0 was used to calculate Mann-Whitney U tests, Kruskal-Wallis tests with p-values adjusted for multiple comparisons via Dunn’s multiple comparison test, and geometric means and geometric standard deviations.

