## Supplementary Figures for "Elicitation of neutralizing antibody responses to HIV-1 immunization with nanoparticle vaccine platforms"

**Supplementary Figure 1. Size-Exclusion Chromatography Profiles of Trimer and Nanoparticle Immunogens.**

Soluble gp140 trimer immunogens were purified on a Superdex 200 Increase 10/300 GL, and nanoparticle immunogens were purified on 16/60 Sephacryl s500 or Superose 6 columns. Dashed lines denote selected fractions.

**Supplementary Figure 2. Comparison of AUC values from ELISA Binding Curves of Guinea Pig Serum Against HIV-1 Env Proteins.**

**A.** AUC values for vaccine-matched Env trimers plotted from ELISA binding curves in Figure 4B**.**

**B.** AUC values for gp140 Env proteins plotted from ELISA binding curves in Figure 4C. Reported are p-values from Mann-Whitney U tests.

**C.** Binding for positive control antibody VRC01 against HIV-1 antigens.

### Supplementary Figure 1

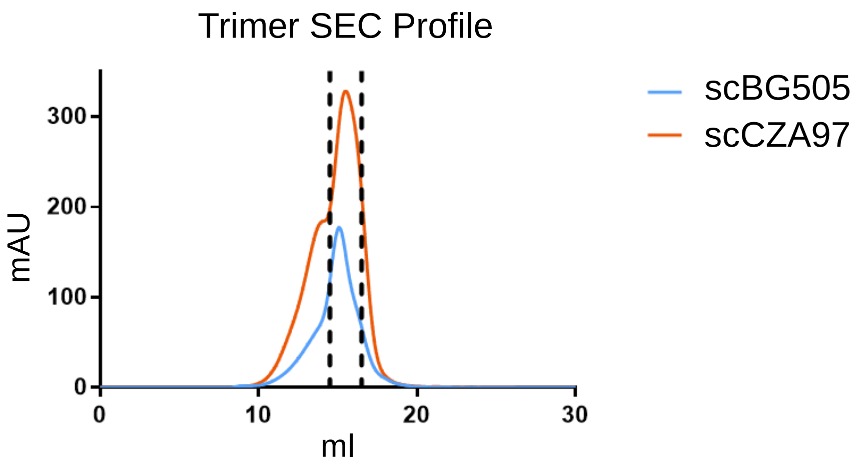


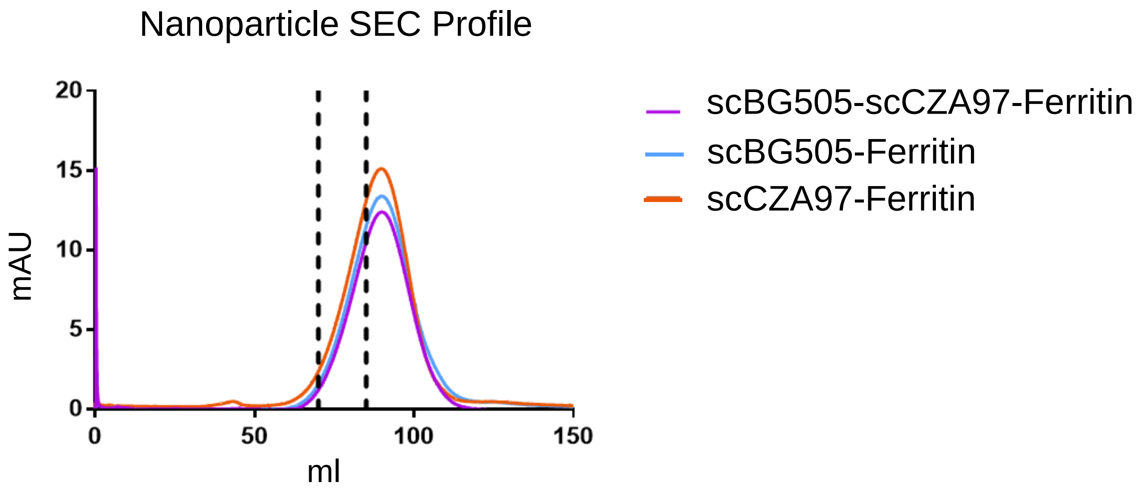

Supplementary Figure 2

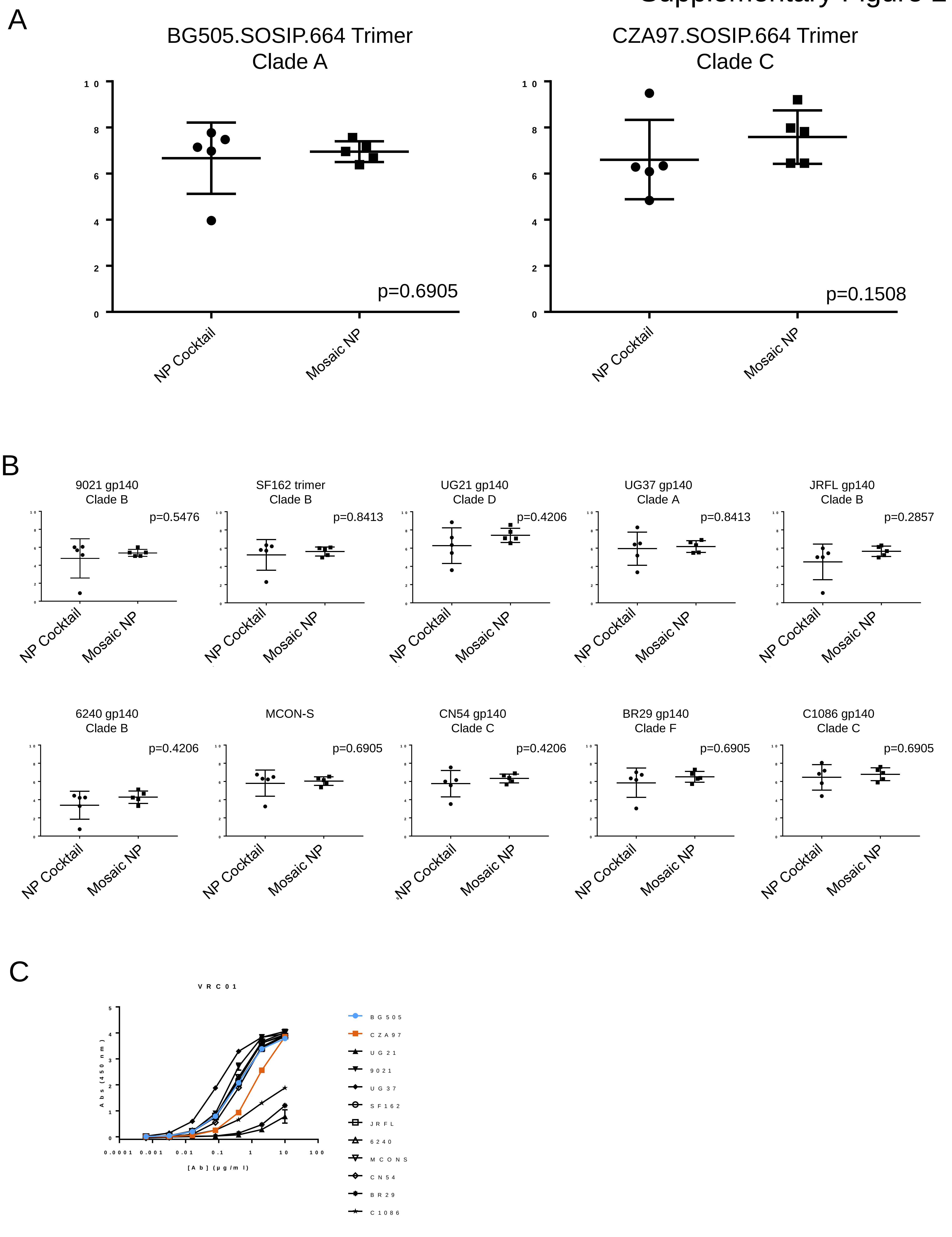
